# DSNetwork: An integrative approach to visualize predictions of variants’ deleteriousness

**DOI:** 10.1101/526335

**Authors:** Audrey Lemaçon, Marie-Pier Scott-Boyer, Penny Soucy, Régis Ongaro-Carcy, Jacques Simard, Arnaud Droit

## Abstract

One of the most challenging tasks of the post-genome-wide association studies (GWAS) research era is the identification of functional variants among those associated with a trait for an observed GWAS signal. Several methods have been developed to evaluate the potential functional implications of genetic variants. Each of these tools has its own scoring system which forces users to become acquainted with each approach to interpret their results. From an awareness of the amount of work needed to analyze and integrate results for a single locus, we proposed a flexible and versatile approach designed to help the prioritization of variants by aggregating the predictions of their potential functional implications. This approach has been made available through a web interface called DSNetwork which acts as a single-point of entry to almost 60 reference predictors for both coding and non-coding variants and displays predictions in an easy-to-interpret visualization. We confirmed the usefulness of our methodology by successfully identifying functional variants in four breast cancer susceptibility loci. DSNetwork is an integrative web application implemented through the Shiny framework and available at: http://romix.genome.ulaval.ca/dsnetwork.

**Author summary:** Over the past years, GWAS have enabled the identification of numerous susceptibility loci associated with complex traits (https://www.ebi.ac.uk/gwas/). However, many of those signals contain hundreds or even thousands of significantly associated variants among which only a few are really responsible of the phenotype. Substantial efforts have been made in the development of prediction methods to prioritize variants within GWAS-associated regions to go from statistical associations, to the identification of functional variants modulating gene expression, in order to ultimately gain insight into disease pathophysiology. Unfortunately, these numerous prediction tools generate contradictory predictions rendering the interpretation of results challenging. Some tools such as VEP [McLaren et al., 2016] report their scores using a color scheme, thus acknowledging the need to assist the user in the interpretation of predictor results. Nonetheless, the multiplication of approaches can often result in an extensive amount of data that is hard to synthesize. Aware of the challenge of evaluating the potential deleteriousness of variants in the context of fine mapping analyses, we created a customizable visualization approach that was implemented it in the decision support tool called DSNetwork for **D**ecision **S**upport **Network**. This tool enables quick access to gold standard and new predictors for both coding and non-coding variants through an easily interpretable visualization of these predictions for a set of variants.

## Introduction

Since 2006, thousands of susceptibility loci have been identified through genome-wide association studies (GWAS) for numerous traits and complex diseases, including breast cancer [MacArthur et al., 2017]. GWAS build on the concept of linkage disequilibrium (LD) to identify statistical associations between genetic variants and diseases [Visscher et al., 2017]. While this approach is powerful for locus discovery, it cannot distinguish between truly causal variants and non-functional highly correlated neighboring variants. Thus, for the vast majority of these loci, the causal variant(s) and their functional mechanisms have not yet been elucidated.

Statistical fine-mapping analyses combined with the functional annotation of genetic variants can help pinpoint the genetic variant (or variants) responsible for complex traits, or at least narrow down the number of variants underlying the observed association for further functional studies. In this regard, tremendous efforts have been put forth to assist the functional assessment of variants at risk loci and numerous scoring methods and tools have been developed to predict the deleteriousness of variants based on a number of characteristics such as sequence conservation, characteristics of amino acid substitution, and location of the variant within protein domains or 3-dimensional protein structure.

In recent years, efforts have been made towards the aggregation of many different functional annotations resulting from these scoring methods in a single integrative value called metascore [Feng, 2017, Ionita-Laza et al., 2016], an approach which seems to yield better performances than any predictor individually [Dong et al., 2015]. Although these methods demonstrate themselves to be useful, they have some limitations, notably they are not directly comparable to each other and their prediction results are sometimes contradictory.

In order to allow a quick survey of a wide range of predictors for a given list of variants and assist in the interpretation of the resulting prediction scores, we propose a flexible and integrative method capable of gathering information from multiple sources in an easy-to-interpret representation rather than a static new metascore. For this purpose, we created a single-point of entry fetching predictors for coding and non-coding variants and presenting them as a network, where the nodes represent the variants of interest and the edges the linkage disequilibrium between variants. The network is built with the aim of rendering the predictor results easier to peruse during analyses involving multiple variants, and therefore, assist in the variant prioritization process in the context of fine-mapping analyses.

This approach has been made available through a web interface called DSNetwork at: http://romix.genome.ulaval.ca/dsnetwork.

## Materials and methods

### Annotations retrieval

Variant annotations and scoring data are fetched on-the-fly from MyVariant.info high-performance web services [Xin et al., 2016] using their third-party R package. SNPnexus [Dayem Ullah et al., 2018] scorings are fetched upon request through a Python script kindly provided by the SNPnexus team. Due to their novelty and relevance for our purpose, three complementary whole-genome resources are included : LINSIGHT [Huang et al., 2017], BayesDel [Feng, 2017] and predictions and sequence constraint data [di Iulio et al., 2018] which can be used as a proxy to score functionality and the consequences of mutations. BayesDel, LINSIGHT and Context-Dependent Tolerance scores were extracted from a local copy. LD data are computed from 1000 Genomes Phase 3 [1000 Genomes Project Consortium et al., 2015].

### Visual integration

Once fetched for the variants of interest, prediction results are displayed as a network, whose components, namely the edges and nodes, are used to convey different types of information in an easy-to-comprehend way.

Nodes correspond to annotated variants and their color scheme displays prediction scores as a pie chart, where each slice represents the score of a variant for a particular predictor. For each predictor, the selected variants are ranked according to their deleteriousness and the rankings are reflected using a color gradient, ranging from green to red, where a red slice indicates a variant more likely to be damaging with regard to a particular predictor. The edges between the nodes can be used to map Linkage Disequilibrium (LD) levels between two variants. LD (squared correlation *r*^2^) is based on a user-chosen 1000 genomes population and is represented by an absolute color gradient ranging from yellow to red. Red indicates a high disequilibrium.

In addition to standard individual predictors, our approach includes overall measures called “metascores”. We provided two types of metascores. The first type consists in an average ranking of all selected predictors, which enables a quick visualization of the global ranking of variants across a particular region. The second type of metascore incorporated several existing integrative approaches namely BayesDel, LINSIGHT, Eigen, Eigen-PC [Ionita-Laza et al., 2016] and an integrative weighted scoring for variations in the noncoding genome called IW-Scoring. This type of metascore provides absolute values comparable between analyses and gives insight into the relevance of each variant regardless of the other candidates. Figure 1 illustrates some networks related to the case studies which will be detailed hereafter.

**Fig 1.**
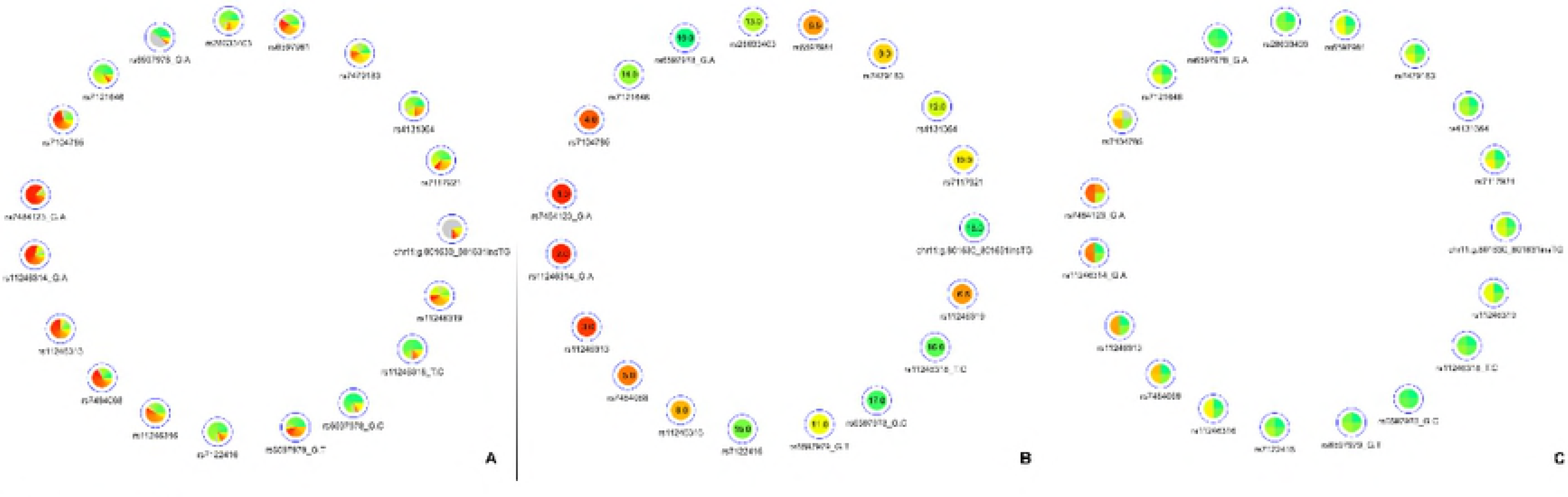
Networks displaying impact prediction scores for a subset of variants of interest. A) network representing all the available predictions; B) relative metascores with missing values ranked at the end; C) absolute metascores showing rs7484123 as the best candidate with regards to deleteriousness predictions.

### Implementation

The DSNetwork was created using the Shiny framework [Chang et al., 2017]. This tool provides the users with deleteriousness predictions for a selected set of coding and non-coding human variants (hg19 build) and generates a set of prioritized results for further analysis. These prediction scores are recovered from several trusted sources and presented in a user-friendly web interface. The interface is organized in three sections, namely Input, Selection and Visualization, as illustrated and described in Figure 2. For complete user guide, see S1 File.

**Fig 2.**
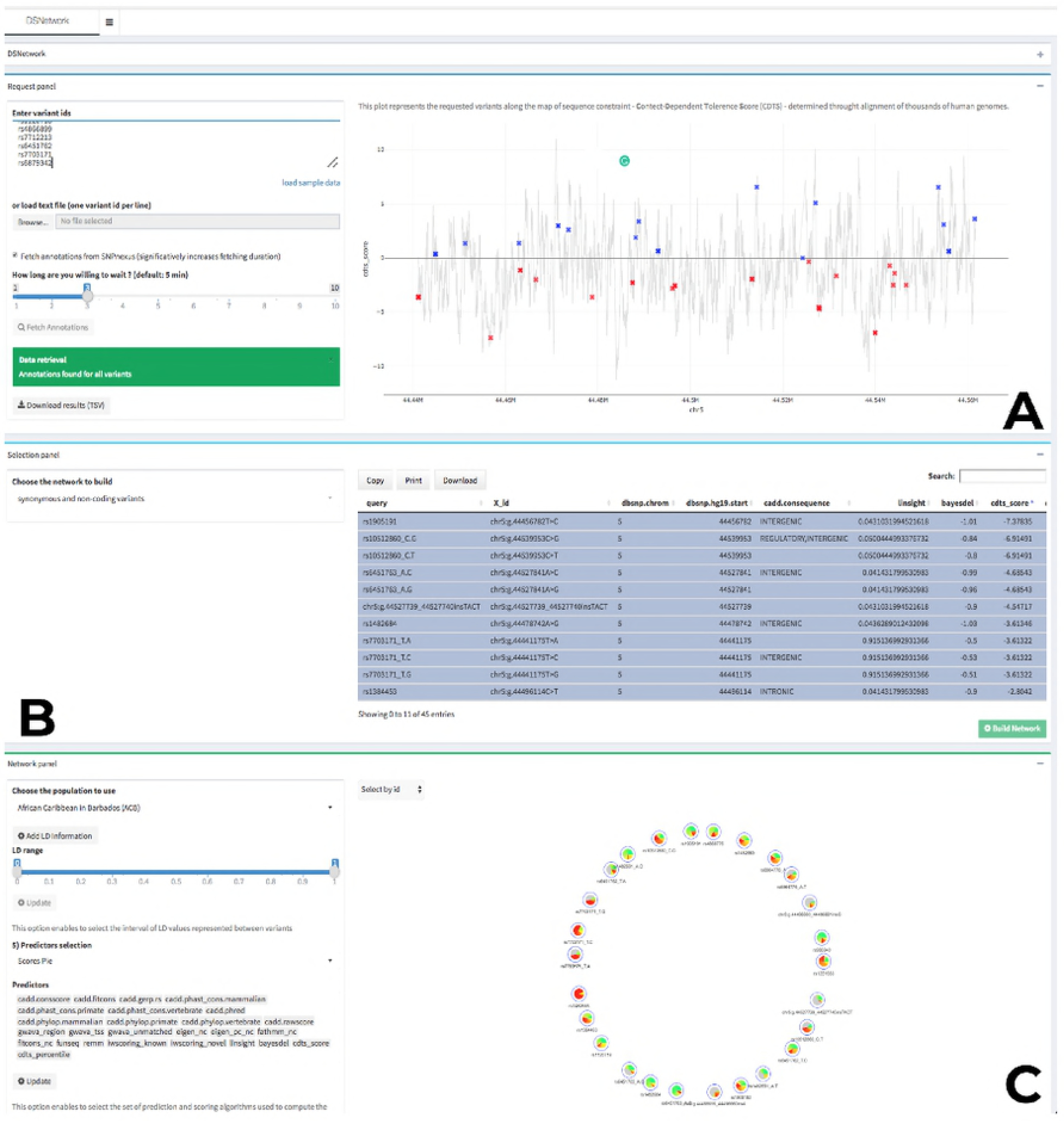
Architecture overview. The first panel is dedicated to user input and parameters for data retrieval. The middle panel presents a relevant subset of annotations for each submitted variant and enables the selection of variants to be integrated in the final visualization. The bottom part on the interface is dedicated to the integrated visualization of the deleteriousness predictions under the form of a network.

### Case studies

We chose to demonstrate the utility of DSNetwork in the context of the functional analysis of four breast cancer susceptibility loci identified through the latest published breast cancer association study (full description in [Michailidou et al., 2017]). This paper reports the discovery of 65 new breast cancer risk loci and deepens the functional characterisation for four regions namely 1p36, 1p34, 7q22 and 11p15. For each of these regions, the authors defined sets of credible risk variants and investigated their impact through functional assays in order to to identify the functional variants.

## Results and Discussion

### Prioritization of four breast cancer susceptibility loci

The original analysis defined for each of the 65 regions, a set of credible risk variants (CRV) containing variants with P-values within two orders of magnitude of the most significant SNPs in this region. They selected four loci for further evaluation namely 1p36, 1p34, 7q22 and 11p15. Initially, those regions contained respectively 54, 13, 19 and 85 significantly associated variants. The p-value cut off enabled to reduce the number of variants to respectively 1, 4, 6 and 19 CRVs. The list of variants for these loci was extracted from the original paper’s Supplementary Tables 8 and 13 in the context the present analysis. Following data extraction, the analysis procedure was: 1) upload the variants of interest on the web tool, 2) fetch the annotations, 3) Visualise the variants through the overview plot, 4) visualise the available deleteriousness scores in the decision network, 5) use relative metascore visualizations to quickly identified the best candidates and finally 6) conclude.

### Locus 1p36

This region contains a single CRV, rs2992756 (P = 1.6*x*10^−15^). For demonstration purposes, we selected the 30 most associated variants to put to the test. Amongst these 30 variants, 2 variants (rs200439143, rs71018084) weren’t annotated by DSNetwork because of their absence from MyVariant.info service, 24 were identified as regulatory variants and 4 as non-synonymous variants. We focused our analysis on the regulatory variants.

Based on the deleteriousness scores available for this subset of variants, a quick overview of variant nodes has allowed to easily identify rs2992756 as the best candidate. Indeed, the node for this variant contained the largest proportion of red, indicating a good ranking for most of the scoring approaches (Figure 3.A). To confirm this observation, we used the relative metascore visualization (Figure 3.B). The mean rankings, clearly materialized by both the color code and the values, enabled the confirmation of rs2992756 as best candidate among the 30 most breast cancer-associated variants at the 1p36 locus. Using reporter assays, Michailidou et al. [Michailidou et al., 2017] demonstrated that the presence of the risk T-allele of this variant within *KLHDC7A* promoter significantly lowers its activity.

**Fig 3.**
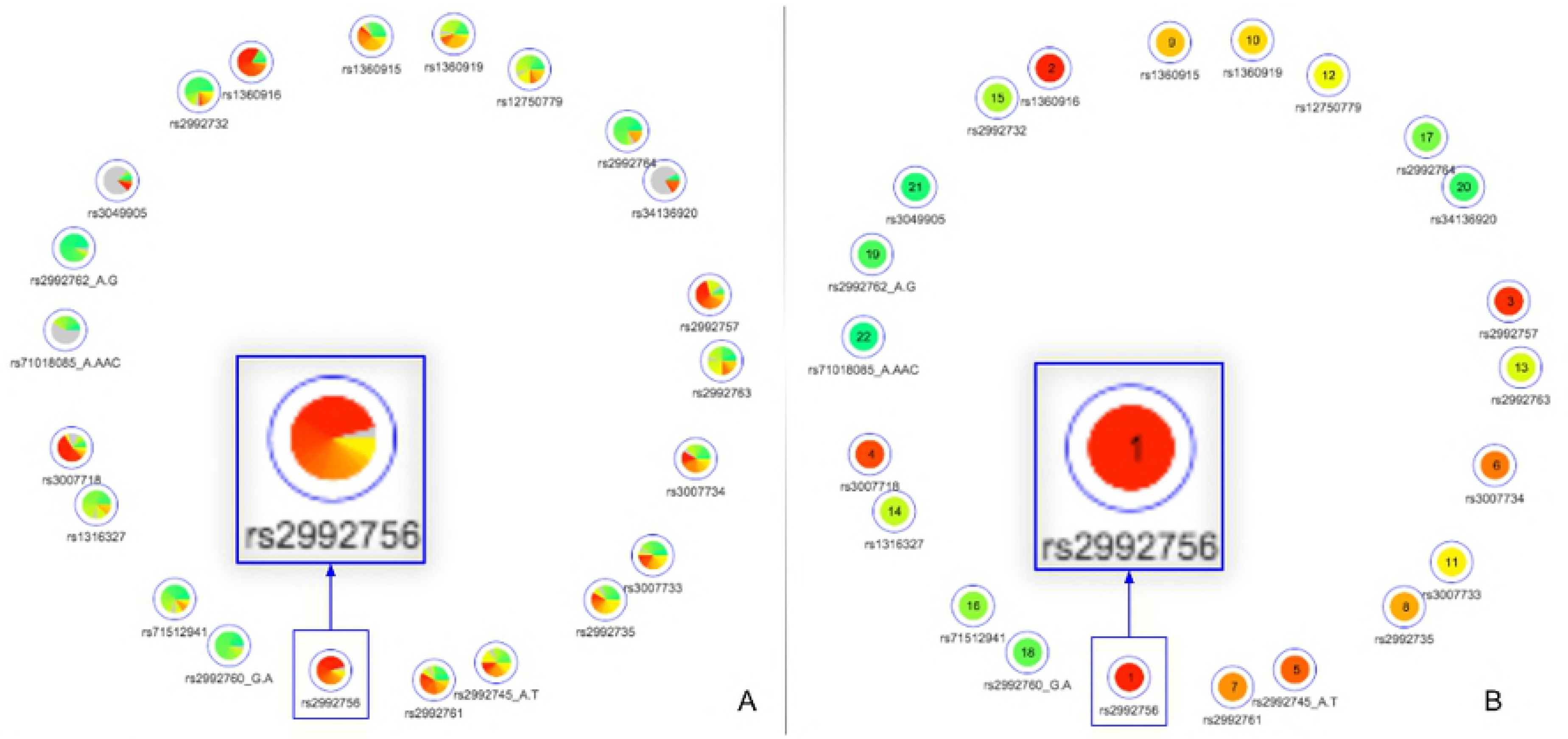
Prioritization analysis of locus 1p36. Networks representing for the 4 CRVs associated variants with breast cancer at the 1p34 locus. A) all available predictions; B) relative metascore with missing values ranked at the end.

### Locus 1p34

This region contains 4 CRVs among 13 significantly associated variants. All the variants were found by DSNetwork and identified as regulatory variants. Based on the deleteriousness scores available for this subset of variants, a quick overview of variant nodes has allowed to easily identify two variants, rs42334486 and rs7554973 as the best candidates. Indeed, the nodes for these variants contained the largest proportion of red, indicating a good ranking for most of the scoring approaches (Figure 4.A). The visualization of the mean ranking confirms rs4233486 as the most credible candidate among the CRVs (Figure 4.B). This observation is in accordance with results from Michailidou et al. [Michailidou et al., 2017], which demonstrated, using reporter assays, that the presence of the risk T-allele of this variant within a putative regulatory element (PRE) reduce *CITED4* promoter activity.

**Fig 4.**
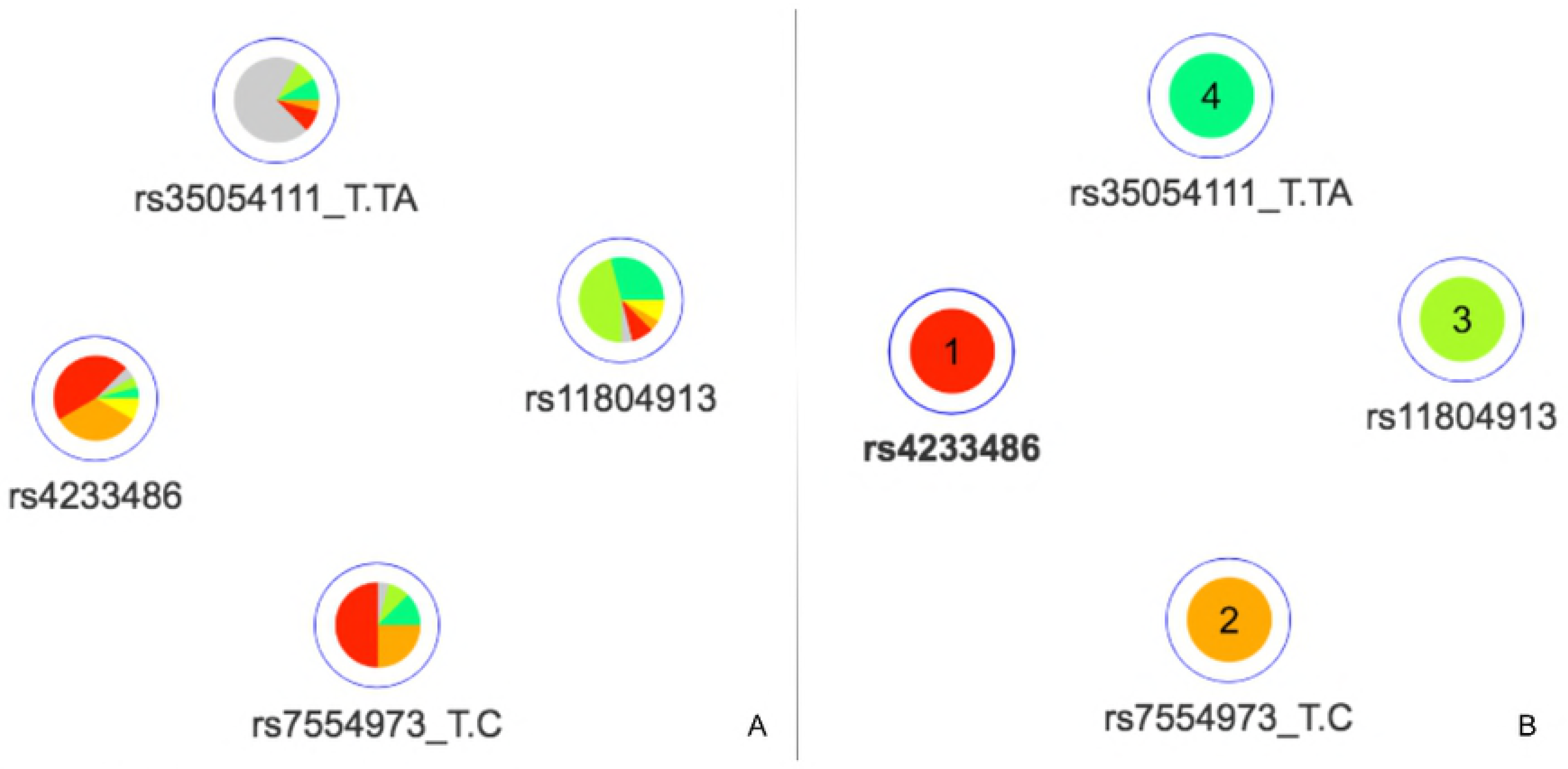
Prioritization analysis of locus 1p34. Networks representing for the 4 CRVs associated variants with breast cancer at the 1p34 locus. A) all available predictions; B) relative metascore with missing values ranked at the end.

### Locus 7q22

This region contains 6 CRVs among 19 significantly associated variants. All the variants were found by DSNetwork and identified as regulatory variants. Based on the deleteriousness scores available for this subset of variants, a quick overview of variant nodes has allowed to easily identify two variants, rs6961094 and rs71559437 as the best candidates. Indeed, the nodes for these variant contained the largest proportion of red, indicating a good ranking for most of the scoring approaches (Figure 5.A). The visualization of the mean ranking confirms rs6961094 and rs71559437 as the most credible candidates among the CRVs (Figure 5.B). This observations are supported by the functional experiments performed by Michailidou et al. [Michailidou et al., 2017], which demonstrated, using allele-specific Chromatin conformation capture (3C) assays, that the presence of the risk-haplotype (rs6961094 combined with rs71559437) is associated with chromatin looping between *CUX1*, *RASA4* and *PRKRIP1* promoters suggesting that the protective alleles abrogate this phenomenon.

**Fig 5.**
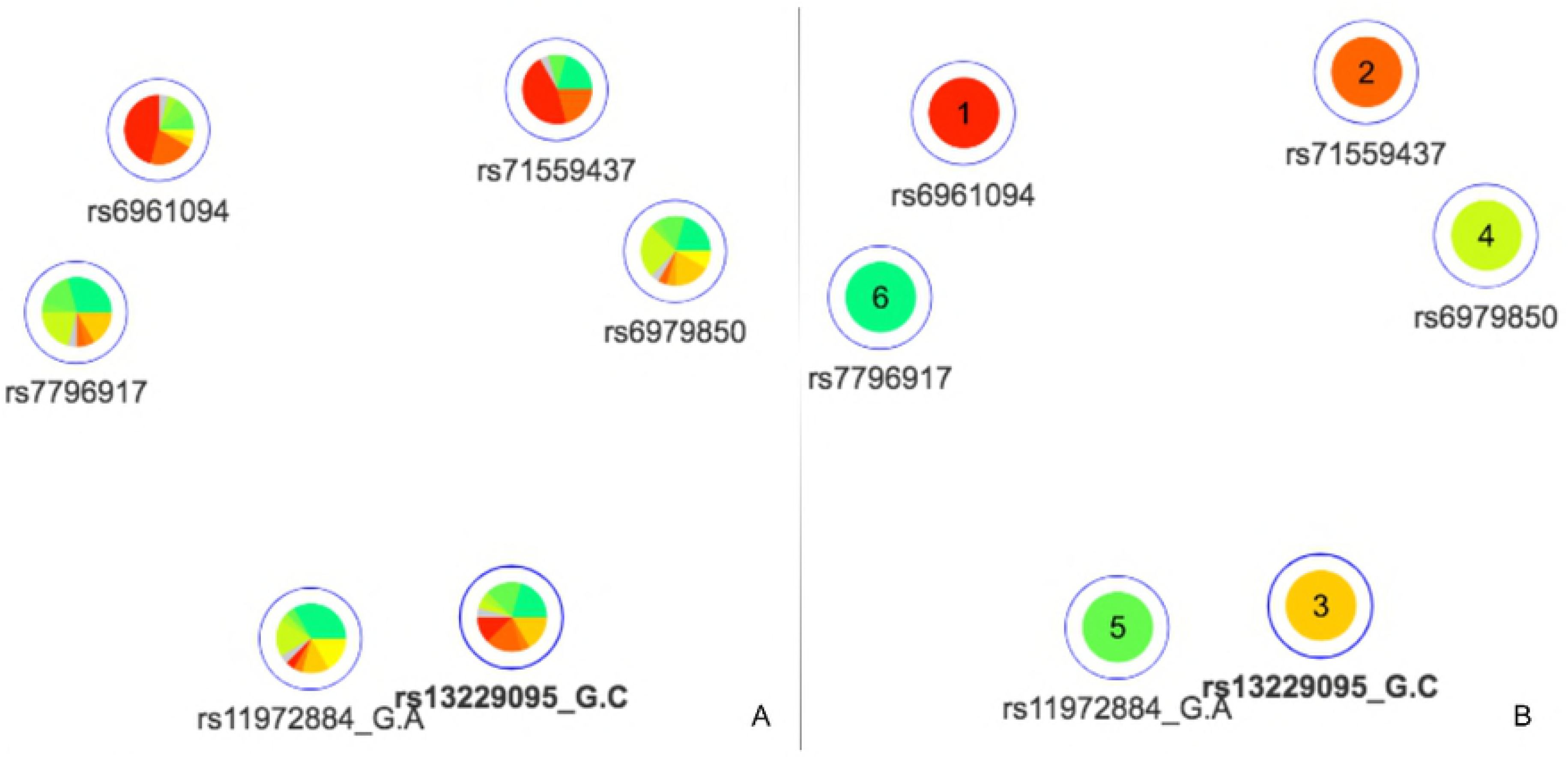
Prioritization analysis of locus 7q22. Networks representing for the 19 CRVs associated variants with breast cancer at the 11p15 locus. A) relative metascore with missing values ranked at the end; B) absolute metascores show rs7484123 and rs11246314 as the best candidates with regard to deleteriousness predictions.

### Locus 11p15

This region contains 19 CRVs among 85 candidate variants. All the variants were found by DSNetwork and 18 were identified as regulatory variants and 1 as a non-synonymous variant. Among the 19 CRVs, five variants, located in the proximal promoter of PIDD1 (a gene implicated in DNA-damage-induced apoptosis and tumorigenesis [Lin et al., 2000], namely rs7484123, rs7484068, rs11246313, rs11246314, rs11246316 were further analysed by [Michailidou et al., 2017]. They demonstrated, using reporter assays, that these variants, incorporated in a construct, significantly increased *PIDD1* promoter activity.

A quick overview of the relative and absolute metascores visualization allowed to easily prioritize the 19 CRVs (Figure 6 A and B). First, the prioritised list based on the metascores confirms the selection of these five variants as functional credible SNPs. Indeed they are ranked at the first, second, third, fifth and eighth place out of nineteen. Moreover, we notice that variants rs7484123 and rs11246314 demonstrate a higher level of coloration, confirming them as the best candidates among the variants located in the proximal promoter of *PIDD1*. The variant rs7484123 particularly stands out as a very promising candidate for subsequent experiments.

**Fig 6.**
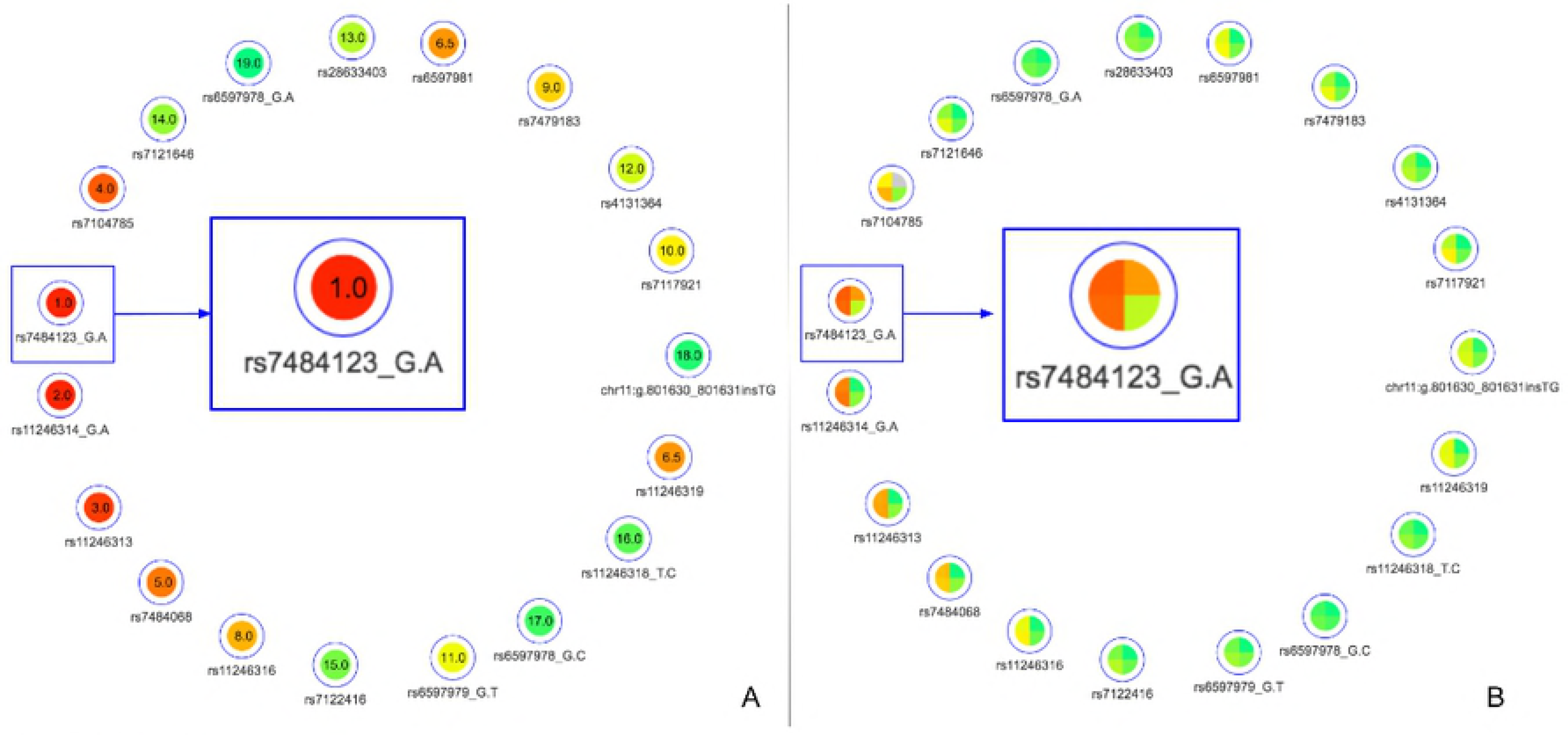
Prioritization analysis of locus 11p15. Networks representing for the 19 CRVs associated variants with breast cancer at the 11p15 locus. A) relative metascore with missing values ranked at the end; B) absolute metascores show rs7484123 and rs11246314 as the best candidates with regard to deleteriousness predictions.

## Conclusion

We analysed the four regions through DSNetwork and were able to pinpoint the same most plausible causal variants than those proposed in the original paper. DSNetwork provides a user-friendly interface integrating predictors for both coding and non-coding variants in an easy-to-interpret visualization to assist the prioritization process. The use of DSNetwork greatly facilitates the selection process by aggregating the results of nearly sixty prediction approaches and easily highlights the best candidate variants for further functional analysis.

## Supporting information

**S1 File.** User guide

## Acknowledgments

We thank the BCAC for providing the impetus to create DSNetwork. We thank the SNPnexus team (Barts Cancer Institute, Queen Mary University of London) for their assistance in integrating SNPnexus data. We also thank Pr. Bing-Jian Feng (University of Utah School of Medicine) for his help and advice regarding BayesDel integration. We are also grateful to all the personnel of Arnaud Droit’s lab and particularly Gwenaëlle Lemoine and Benjamin Vittrant for their advice and assistance in the preparation of this article. We wish to extend our thanks to all the current and future DSNetwork users.

